# Kinetic model derivation for design, building and operation of solid waste treatment unit based on system dynamics and computer simulation

**DOI:** 10.1101/2022.11.18.516877

**Authors:** Dawei Hu, Shuaishuai Li, Xiaolei Liu, Hong Liu, Guanghui Liu

## Abstract

Solid waste disposal is significantly important to maintain normal operation of both natural and artificial ecosystems. In this study, a kinetic model of solid waste treatment unit (SWTU) was upfront developed based on microbial ecology, system dynamics, cybernetics and digital simulation, which accurately described the relationships and interactions between solid waste decomposition (SWD) processes and biotic/abiotic factors. Then a specific SWTU prototype was designed and built from this kinetic model. A 370-day experiment demonstrated that SWTU maintained normal operation with robust stability and desired dynamic behaviors, and effectively disposed the solid waste. Therefore, this kinetic model was highly valid due to its high structural and behavioral similarity with the prototype. This research could lay a strong theoretical foundation for further closed-loop control and optimization of SWTU, and provide scientific guidance to environmental management and sustainable development.

## 1. Introduction

Solid waste, such as plant debris, animal excrement, must be decomposed and utilized by microorganisms, releasing biogenic materials back into the environment for the next ecological circulation and ensuring the perpetual operation of the natural ecosystem(Hamer, 2003). In order to emulate this natural ecological process, various solid waste decomposition technologies have been currently developed for engineering application and played a vital role in environmental protection, ecological restoration(Lim et al., 2016), and artificial ecosystem construction(G. Liu et al., 2016).

Nevertheless, spontaneous SWD processes are relatively slow and unsatisfying for engineering application. Therefore, it is currently vital to design and build an extremely-efficient solid waste treatment unit to accelerate SWD processes based on dynamic mechanisms of SWD—namely the interactions and relationships between SWD and biotic/abiotic environmental factors.

However, as is known, SWD processes are considerably complicated with nonlinear and time-variant properties, causing their complex dynamic behaviors(D. Liu et al., 2018). Therefore, it is practically impossible to design, build, and operate the SWTU prototype only based on empirical trial-and-error experiments. The model-based design and building for prototype system has been the preferable way to address the problem in information era in a holistic, efficient and economical mode (Ushakova et al., 2012)—The emerging consensus around complex systems science (Nanda & Berruti, 2021; Nordlund et al., 2009) indicates that this maybe the only way to systematically investigate the complicated structure and dynamic behaviors of living systems, especially the profound reciprocal effect and relationships between living systems and environmental factors.

So far, many studies have been conducted to investigate the interactions and relationships between kinetic model of SWD processes and environmental factors. For example, Białobrzewski (2015a) presented a model to describe the downwelling sludge fermentation, whose results showed that temperature affected the fermentation because of different bioactivity among species at the same temperature. Ahn (2008) analyzed 12 types of fermentation material at different water contents and predicted the effect of moisture and compost height on porosity and permeability. Vasiliadou (2015a) predicted the amount of oxygen consumed by the biomass through a model which could lead to the selection of the proper aeration strategy to improve the waste treatment process. Rafiee (2017) developed a mass balance model for estimating the rate of composting and found the law of coexistence of aerobic and anaerobic fermentation. Solemauri (2007) provided an insightful analysis of the three-phase change of matter, making the integrated model more accurate and practical.

Although most of studies focused on the influence of a single or a few environmental factors in SWD processes, under the practical circumstance, the SWTU operation would be affected by all environmental factors together, including microbial biomass, temperature, water, O_2_, minerals, and so on. Furthermore, the relationships between SWD processes and biotic/abiotic factors were often considered one-way in the researches mentioned above, and these researches did not take strong coupling relationships between them into account, i.e., the feedback effect of SWD processes on biotic/abiotic factors into consideration. Therefore, these mathematical models cannot effectively predict the dynamic response characteristics of SWTU to theoretically guide its design, building and operation.

Bioregenerative Life Support System (BLSS) is a specific closed artificial ecosystem composed of human-beings, higher plants, insects, microorganisms and artificial environments (including light, temperature, water, gases, mineral substances, and so on). “Lunar Palace 1 (LP1)” is constructed based on it and will be applied for manned space exploration in the future(G. Liu et al., 2016). A 370-day experiment with four crew members was successfully performed in LP1, where mass of solid waste (such as inedible biomasses of higher plants, crew members waste, insects frass, kitchen garbage) was continuously produced during the experiment(Fu et al., 2016). Because the rate of solid waste disposal restricts the total mass transportation and transformation, it is considered the “bottleneck” of matter turnover in the field of BLSS. Therefore, how to design and build the extremely efficient SWTU has become a tremendously pressing problem for the successful operation of BLSS.

In this article, based on microbial ecology, system dynamics, cybernetics, combined with previous researches mentioned above, a kinetic model describing the inner structure of SWTU (i.e., the interactions and relationships between SWD processes and biotic/abiotic factors) was upfront developed to elucidate the complicated dynamic mechanisms. These mechanisms were implemented to drive the SWTU operation and obtain dynamic responses to variations of environmental factors through numerical simulation. Then the model-based design and building of SWTU prototype was performed. The 370-day experiment proved the SWTU operated with high reliability as well as stability, and effectively disposed the solid waste to guarantee the normal operation of LP1. The parameter identification and model validation were conducted by experimental data, which proved the kinetic model and prototype of SWTU were exceedingly similar in structure and dynamic behaviors.

To our knowledge, none of the existing SWD process models has comprehensively taken microbial biomass, inedible biomass, fermentation material temperature, water content in fermentation material, O_2_ concentration in fermentation material as well as their strong coupling relationships into account at the same time. Therefore, the objective of this study is to develop a highly valid kinetic model with the strong coupling of SWD processes and key biotic/abiotic factors. Special emphasis has been placed on design, building and operation of SWTU prototype based on kinetic model and prediction of its dynamic behavior performances.

## 2. Materials and methods

### 2.1 Kinetic model derivation

#### 2.1.1 Hypotheses and the flow chart in modeling SWTU

In order to deeply investigate the complicated dynamic characteristics of the SWTU, a grey-box model expressed by a set of parameterized nonlinear first-order differential equations (ODEs) is developed based on related mechanisms, experimental data and system dynamics, according to the following fundamental assumptions:

1. The components of SWTU only change with time, which means space is not taken into account. Hence, its system dynamic model with a lumped parameter can be described by ODEs.
2. The primary kinetic process in the SWTU is the microbial growth, and the rates of other non-living processes pertinent to it were obtained correspondingly by the coefficient conversion method.
3. SWTU operates in the normal state, signifying the structure and function of the microbial community and abiotic factors are all in the normal state during SWTU operation.
4. Although there are many species in the aerobic microbial community, their interspecific differences are ignored for simplification of model derivation, namely it is believed that microorganisms have the same growth and metabolic characteristics.
5. Although solid waste is composed of higher plants inedible biomass, crew members waste, insect’s frass and kitchen garbage, the first takes overwhelming majority and is more difficult to be decomposed in contrast with the others. Therefore, only inedible biomass is taken into consideration.
6. SWTU is a time-invariant system, because parameters of the kinetic model could be considered unchangeable during operation.

According to hypotheses, aerobic fermentation process mechanisms and design of four-phase material transport, the flow chart of inner processes in SWTU (Fig.1) could be obtained correspondingly.

**Fig. 1:**
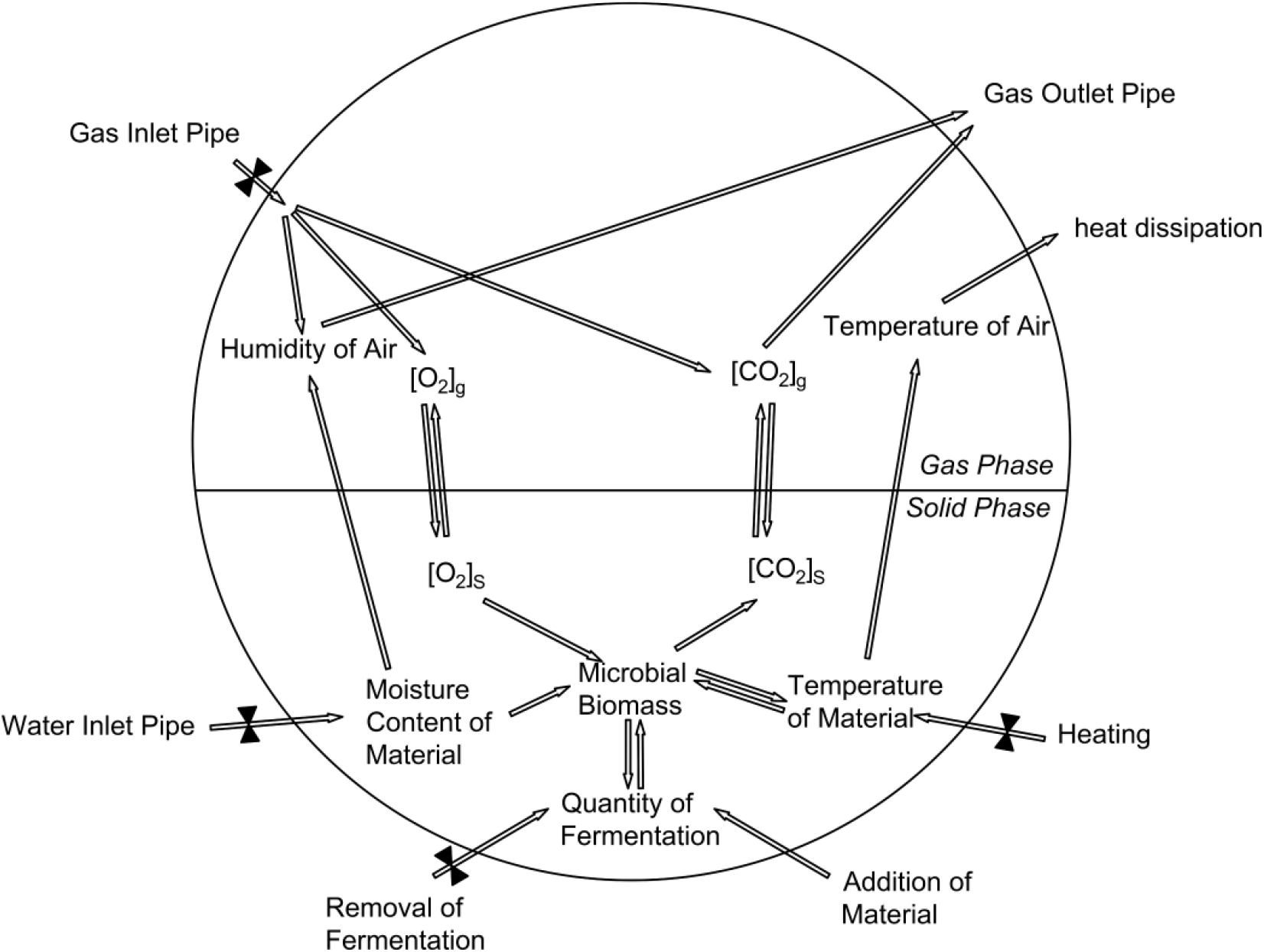
Flow chart of inner processes in SWTU. Based on the flow chart of inner processes in SWTU, the components represented by state variables, relationships and interactions between components are quantitatively described by rate equations with parameters regulating the strength of interconnection and interaction, and a valid kinetic model of SWTU could be developed to accurately express the internal structure and external dynamic behaviors of SWTU.

#### 2.1.2 State variables

As illustrated in Fig. 1, the kinetic model of SWTU was composed of 11 state variables according to the flow chart as follows:

Microbial biomass (*x*_*1*_: g), inedible biomass (*x*_*2*_: g), ash quantity (*x*_*3*_: g), fermentation material temperature (*x*_*4*_: °C), water content in fermentation material (*x*_*5*_: g), O_2_ concentration in fermentation material (*x*_*6*_: %), CO_2_ concentration in fermentation material (*x*_*7*_: %), O_2_ concentration in SWTU gas phase (*x*_*8*_: %), CO_2_ concentration in SWTU gas phase (*x*_*9*_: %), temperature of SWTU gas phase (*x*_*10*_: °C) and humidity of SWTU gas phase (*x*_*11*_: %).

#### 2.1.3 Rate equations

Rate equations were derived to quantitatively express the interactions and relationships driving SWTU operation as follows:

1. Rate equations related to microbial biomass (*x*_*1*_) Microbial growth rate (*v*_*1g*_) was directly affected by microbial biomass (*x*_*1*_), inedible biomass (*x*_*2*_), fermentation material temperature(*x*_*4*_), water content in fermentation material(*x*_*5*_), O_2_ concentration in fermentation material (*x*_*6*_), and could be written as Eq. (1) based on the minimum limiting principle.

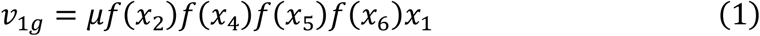

where *μ* denotes the maximum specific growth rate of the microorganisms, *f* (*x*_*2,4,5,6*_) represents the influence function of inedible biomass, fermentation material temperature, water content in fermentation material, O_2_ concentration in fermentation material, respectively. All range between 0 and 1. The influence function of inedible biomass (*x*_*2*_) is formulated by the classical Monod equation as below:

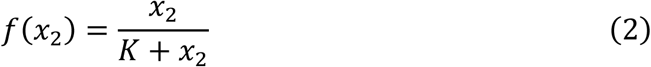

where *K* is the substrate-dependent half-saturation constant of the growth of microorganisms. The influence function of temperature (*x*_*4*_) can be expressed as:

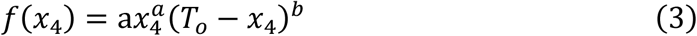

where *T*_*o*_ is the upper limit temperature for growth of microorganisms, *a* and *b* are experimental constants. The *f*(*x*_4_) is also a bell-like function with a necessarily simplified form for engineering application, in contrast to other effective models of temperature influencing microbial growth (Rosso et al., 1995). Similarly, it is harmful to microorganisms to live in too high or too low moisture content (*x*_*5*_), thus the influence function of moisture can be defined as:

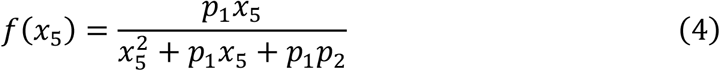

where *p*_*1*_ and *p*_*2*_ are experimental constants. Although both Equations (3) and (4) indicate that high or low levels of environmental factors adversely affect microbial growth, their function shapes are different because Eq. (3) varies fast with *x*_4_, while Eq. (4) varies slowly with *x*_5_. Because the SWD processes are caused by microbial aerobic fermentation, the influence function of O_2_ concentration can be also expressed in the form of Monod function as below:

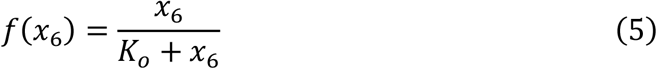

where *K*_*o*_ is the half-saturation constant of the microbial growth on O_2_ concentration. Combining Eq. (1) ∼ (5), the microbial growth rate can be expressed as:

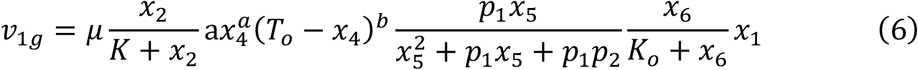

The microbial metabolism rate (*v*_*1d*_) can be expressed as:

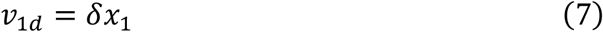

where *δ* is the natural metabolism coefficient of the microorganism. The fermentation material mainly consists of inedible biomass of plants, ash, water and microorganisms, while the gas could be neglected due to relatively lightweight. Therefore, the total mass of fermentation material is:

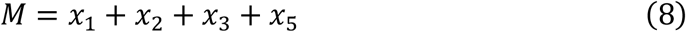

Hence, the discharging rate of microorganisms(*v*_*1o*_) can be expressed as:

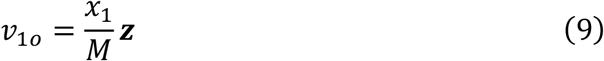

In Eq. (9), ***z*** (Kg h^-1^) denotes discharging speed during the SWTU operation.
2. Rate equations related to inedible biomass (*x*_*2*_) The rate of consumption of substrate by microorganisms during growth(*v*_*2d*_) and discharging of microorganisms(*v*_*2o*_) can be written as follows:

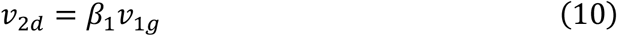

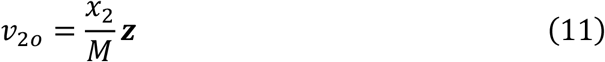

where *β*_*1*_ denotes the microbial consumption coefficient of inedible biomass.
3. Rate equations related to ash quantity (*x*_*3*_) The rate of ash production from substrate decomposition during microbial growth(*v*_*3g*_) and discharge of microorganisms(*v*_*3d*_) could be written as follows:

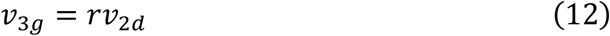

where *r* denotes the proportion of ash element in inedible biomass, and,

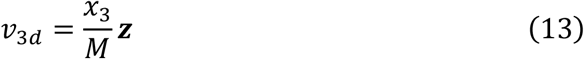
4. Rate equations related to fermentation material temperature(*x*_*4*_) The temperature change follows the first law of thermodynamics, i.e., the law of energy conservation. In this study, the rates of increase(*v*_*4g*_) and decrease(*v*_*4d*_) of fermentation material temperature are established by Fourier’s Law of heat conduction:

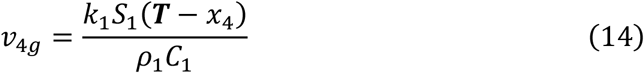

where *k*_*1*_ denotes the heat transfer coefficient of fermentation material, ρ_1_ denotes the density of fermentation material, *C*_*1*_ denotes the heat capacity of fermentation material, *S*_*1*_ is the area of the heat sink, ***T*** is the current temperature of the heat sink. The decrease in fermentation material temperature is due to heat convection and heat radiation into the gas phase.

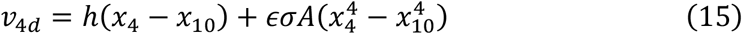

Therefore, the first term on the right in Eq. (15) denotes the thermal convection, where *h* is thermal convection conversion coefficient; the second term denotes thermal radiation, where *ε* is fermentation material emissivity, *σ* is Stefan-Boltzmann constant, and *A* is the area occupied by the material (only considering the bottom area of SWTU).
5. Rate equations related to water content in fermentation material (*x*_*5*_) They include the flow rate of inflow pipe (***Q***) leading to an increase in the water content of fermentation material and water evaporation to the air(*v*_*5d*_). The Priestly-Taylor model is used to express the evaporation rate of water, which has a simple form to be easily calculated as follows:

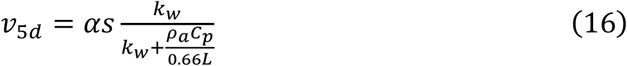

where *α* is the experimental coefficient; *k*_*w*_ is the ratio of vapor pressure in the gas phase and temperature. Both can be determined by linear regression analysis by look-up table; *s* is the heat flux of the fermentation material; *ρ*_*a*_ and *C*_*p*_ are the density and specific heat capacity of air, respectively; and *L* is the evaporation latent heat equivalent of water in the fermentation material. The discharging rate of water (*ν*_50_) was written as follows:

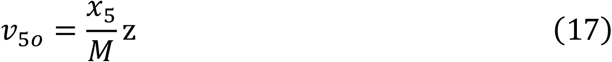
6. Rate equations related to O_2_ concentration in fermentation material(*x*_*6*_) They include the rate of O_2_ permeating into the fermentation material to be mainly dissolved in water from the gas phase (*v*_*6g*_) and rate of O_2_ utilized by microbial respiration(*v*_*6d*_), where O_2_ entering and exiting from the inflow pipe and discharge port is little enough to be neglected. Therefore, *v*_*6g*_ can be expressed by the gas-liquid dual membrane model as below:

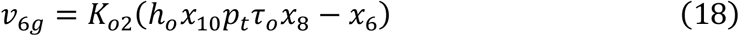

where *K*_*o2*_ is the mass transfer constant of O_2_ between gas and liquid phases, *h*_*o*_*x*_*10*_ is Henry’s constant of O_2_, and *h*_*o*_ represents the regression coefficient between Henry’s constant of O_2_ and temperature, *τ*_*o*_ is the conversion factor of mass and volume of O_2_ in the gas phase, which is determined by the ideal gas law, *p*_*t*_ was the total gas pressure in the gas phase because the sum of nitrogen and water vapor partial pressure was far larger than that of O_2_ and CO_2_, thus *p*_*t*_ could be considered as a constant, *h*_*o*_*x*_10_*p*_*t*_*τ*_*o*_*x*_8_ denotes the saturation concentration of O_2_ in the liquid phase, which is closely related to the concentration and mass transfer process of O_2_. The growth of microorganisms is accompanied by the consumption of O_2_ in fermentation material, therefore the rate equation of *ν*_6*d*_ can be expressed as follows:

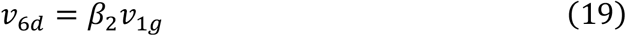

where *β*_*2*_ denotes O_2_ consumption coefficient of microorganisms.
7. Rate equations related to CO_2_ concentration in fermentation material (*x*_*7*_) Similarly, CO_2_ in the fermentation material is also dissolved in wa
ter, to which the rates related include that of CO_2_ permeating from the fermentation material into the gas phase(*v*_*7d*_) and produced by microbial respiration(*v*_*7g*_), while CO_2_ entering and exiting from the inflow pipe and discharge port as well as variation caused by hydrolysis is little enough to be neglected. Therefore, *v*_*7d*_ can be expressed by the gas-liquid dual membrane model as below:

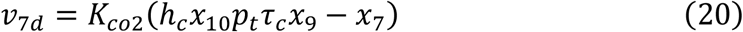

where *K*_*co2*_ is the mass transfer constant of CO_2_ between gas and liquid phases, *h*_*c*_*x*_*10*_ is Henry’s constant of CO_2_, *h*_*c*_ represents the regression coefficient between Henry’s constant of CO_2_ and temperature, *τ*_*c*_ is the conversion factor of mass and volume of CO_2_ in the gas phase, which is also determined by the ideal gas law, and *h*_*c*_*x*_10_*p*_*t*_*τ*_*c*_*x*_9_ denotes saturation concentration of CO_2_ in the liquid phase, which is closely related to the CO_2_ concentration in the gas phase and mass transfer process of CO_2_. The growth of microorganisms is accompanied by the production of CO_2_ in fermentation material, so its rate equation (*ν*_7*g*_) can be expressed as below:

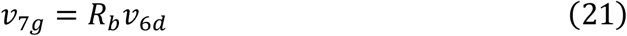

where *R*_*b*_ denotes the respiratory quotient of microorganisms.
8. Rate equations related to O_2_ concentration in SWTU gas phase(*x*_*8*_) The rates of O_2_ entering from the inlet pipe (*v*_*8i*_), permeating from the liquid phase to the gas phase (*v*_*8g*_) and exiting from the outlet pipe (*v*_*8o*_) can be expressed as follows:

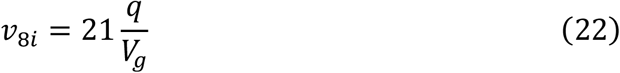

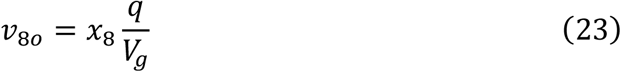

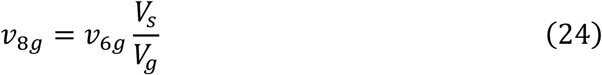

In Eq. (22) ∼ (24), ***q*** is the current flow rate of both the inlet and outlet pipe; 21 is the volume percentage of O_2_ in the outside air; *V*_*g*_ is the volume of the gas phase of the SWTU; *V*_*s*_ is the sum of volume of solid, liquid and biomass phases.
9. Rate equations related to CO_2_ concentration in SWTU gas phase(*x*_*9*_) Similar to O_2_, rates of CO_2_ entering from the inlet pipe (*v*_*9i*_), permeating from the liquid phase to the gas phase (*v*_*9g*_) and exiting from the outlet pipe (*v*_*9o*_) can be expressed as follows:

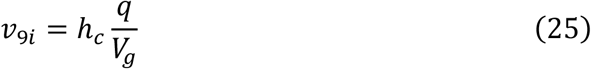

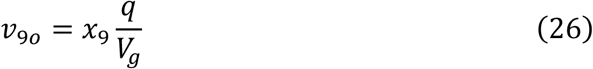

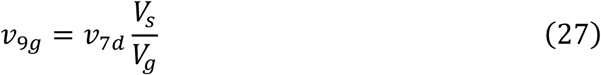

where *h*_*c*_ is the volume percentage of O_2_ in the outside air.
10. Rate equations related to the temperature of SWTU gas phase(*x*_*10*_) The temperature of the air in SWTU increases due to heat convection and heat radiation from fermentation material (*v*_*4d*_), and decreases due to heat released from the metal shell of SWTU to the outside (*v*_*10d*_) expressed as below:

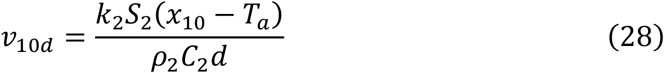

In Equations (28), *k*_*2*_ is the thermal conductivity of the metal wall of SWTU, *ρ*_*2*_ is density of that, *C*_*2*_ is heat capacity of that, *d* is thickness of that, S_2_ is the surface area of the inner shell of SWTU, *T*_*a*_ is the temperature of the external environment of SWTU, which is a certain value. In addition, heat taken away from the outlet pipe is small enough to be neglected.
11. Rate equations related to humidity of SWTU gas phase (*x*_*11*_) They include rate equations of moisture entering from the inlet pipe (*v*_*11i*_), evaporating from the liquid phase to the gas phase (*v*_*11g*_) and exiting to the outlet pipe (*v*_*11o*_).

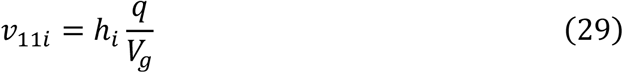

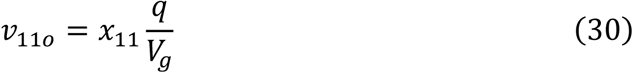

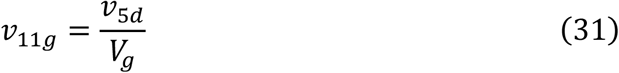

where *h*_*i*_ is the volume percentage of water vapor in the outside air, namely air humidity.

#### 2.1.4 Dynamic model of SWTU

Based on the preceding analysis and modeling process, the kinetic model of SWTU, i.e., a state-space model with 11 first-order nonlinear kinetic equations, could be formulated in a top-down manner as follows:

1. Microbial biomass (*x*_*1*_)

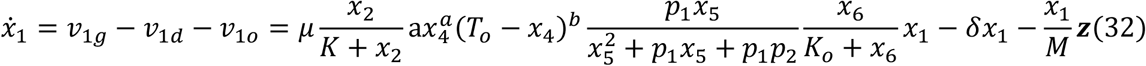
2. Inedible biomass(*x*_*2*_)

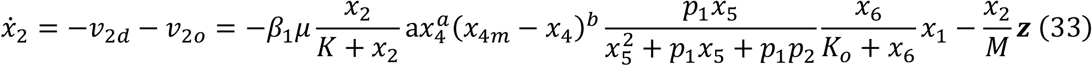
3. Ash quantity(*x*_*3*_)

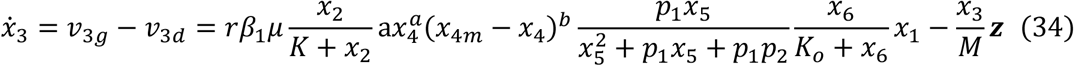
4. Fermentation material temperature(*x*_*4*_)

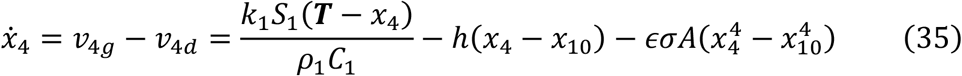
5. Quantity of water in fermentation material(*x*_*5*_)

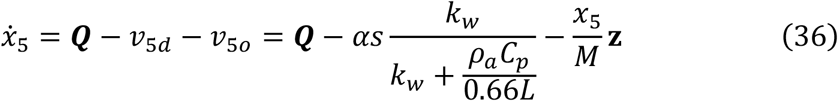
6. O_2_ concentration of fermentation material(*x*_*6*_) in SWTU

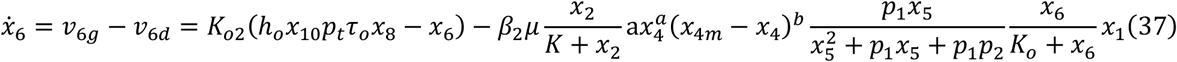
7. CO_2_ concentration of fermentation material in SWTU (*x*_*7*_)

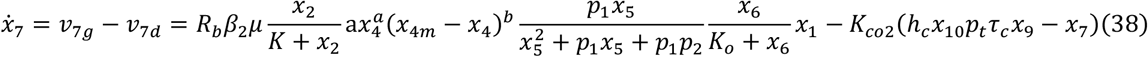
8. O_2_ concentration of gas phase(*x*_*8*_) in SWTU

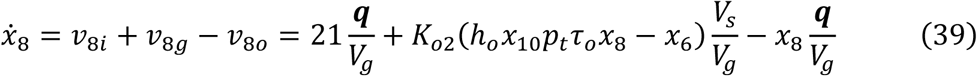
9. CO_2_ concentration of gas phase(*x*_*9*_) in SWTU

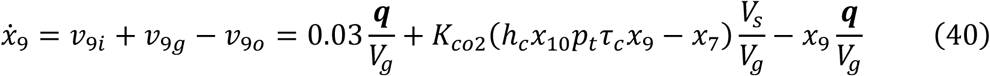
10. Temperature of SWTU gas phase(*x*_*10*_)

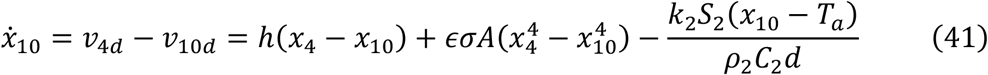
11. Humidity of SWTU gas phase (*x*_*11*_)

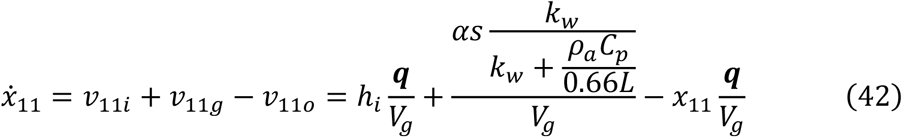

Based on the above SWTU kinetic model (Eq.(32)∼ (42)), the simulation of SWTU can be established for parameter identification and model validation, and further system design, analysis, synthesis, control and optimization by computer simulation.

### 2.2 Simulation model of SWTU

The simulation model of SWTU and its subsystems was built on the MatLab/Simulink platform (Mathworks & Inc, 2016). For example, the simulation model of the microbial subsystem (Fig. 2) is established based on a kinetic model of the microbial population (Eq. 32).

**Fig. 2:**
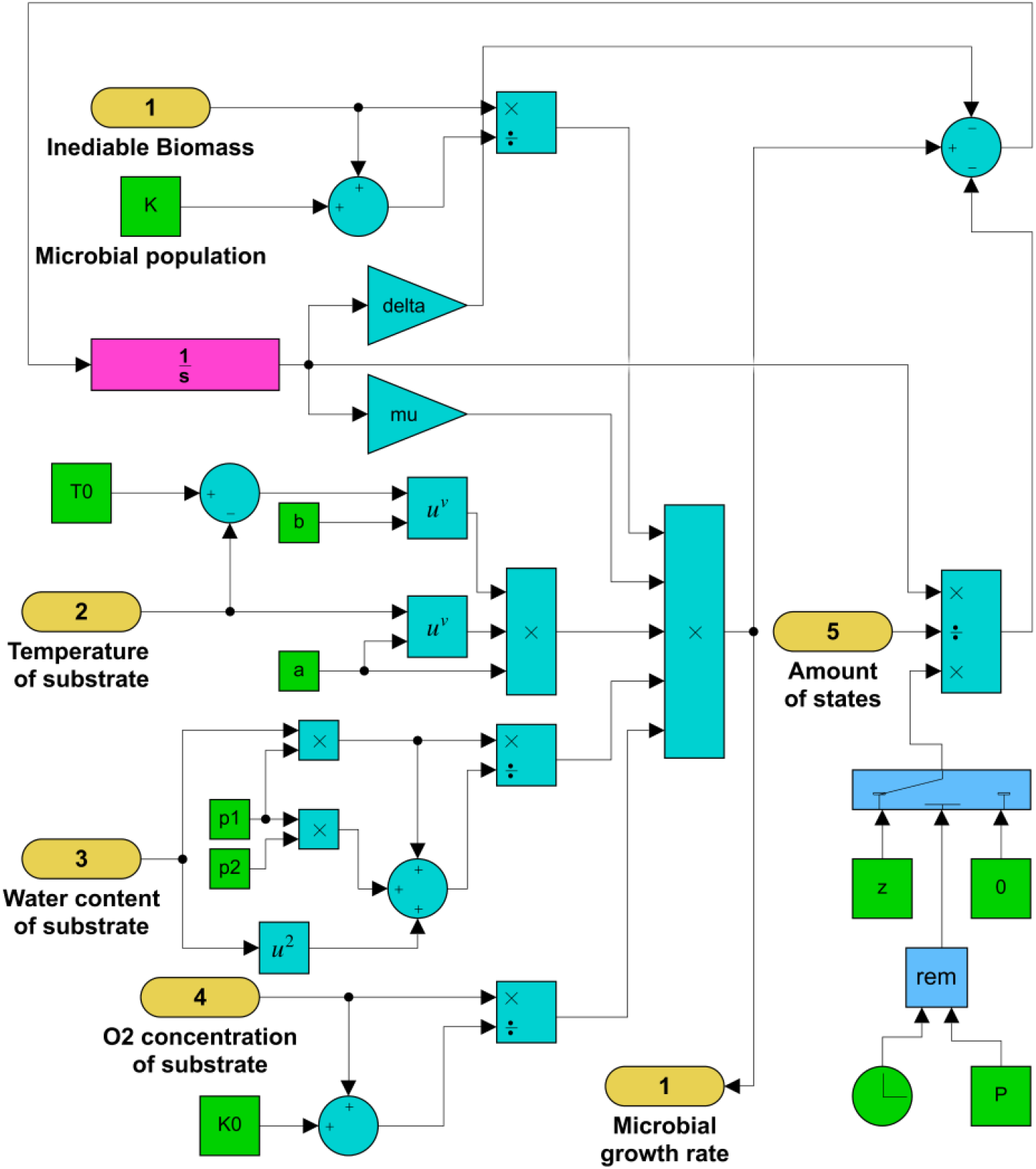
The simulation model of the microbial growth process.

Similarly, the simulation model of the other subsystems can also be respectively established in the same way, and multidimensionally coupled together to form the whole entire SWTU model via input-output ports (Fig. 3).

**Fig. 3:**
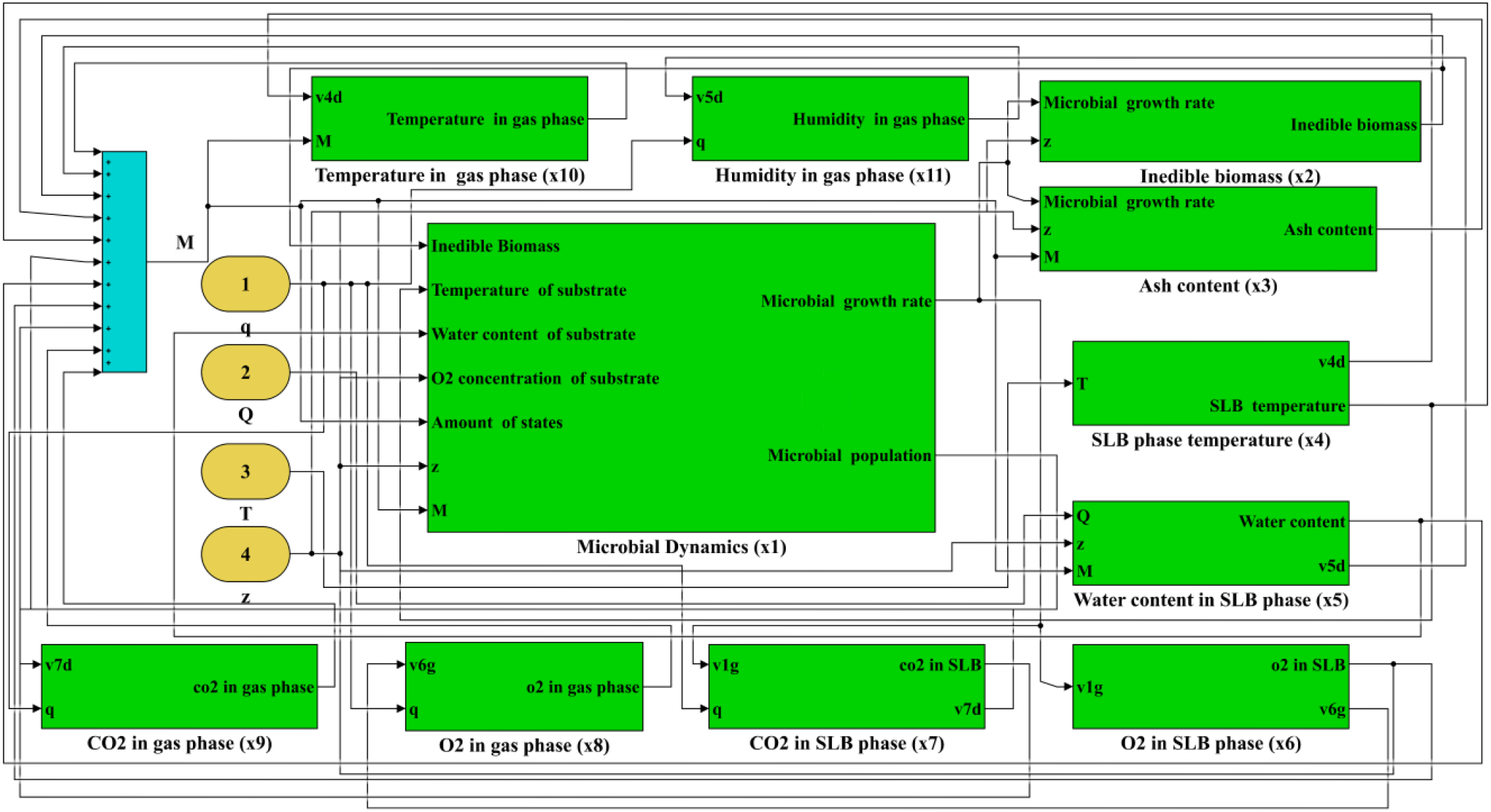
The whole simulation model applied for design and building of SWTU prototype.

Based on the whole simulation model and numerical simulation, some prescribed SWD behavior characteristics with proper delay time, rise time, peak time, settling time, degradation rate, energy consumption and so forth, have been put forward beforehand, then the parameters in the kinetic model (Eq. (32) ∼ (42)) were theoretically specified by dynamic response optimization, therefore a desired digital SWTU was established for guidance of model-based design, building and operation of SWTU prototype. However, the actual parameters might be different from these theoretical parameters and must be identified by experimental data obtained in the course of SWTU prototype operation.

### 2.3 SWTU Prototype

The SWTU prototype, including gas, solid, liquid and biological phases, was designed and built from its kinetic model, and then bottom-up assembled together to obtain the integrated SWTU prototype (Fig.4).

**Fig. 4:**
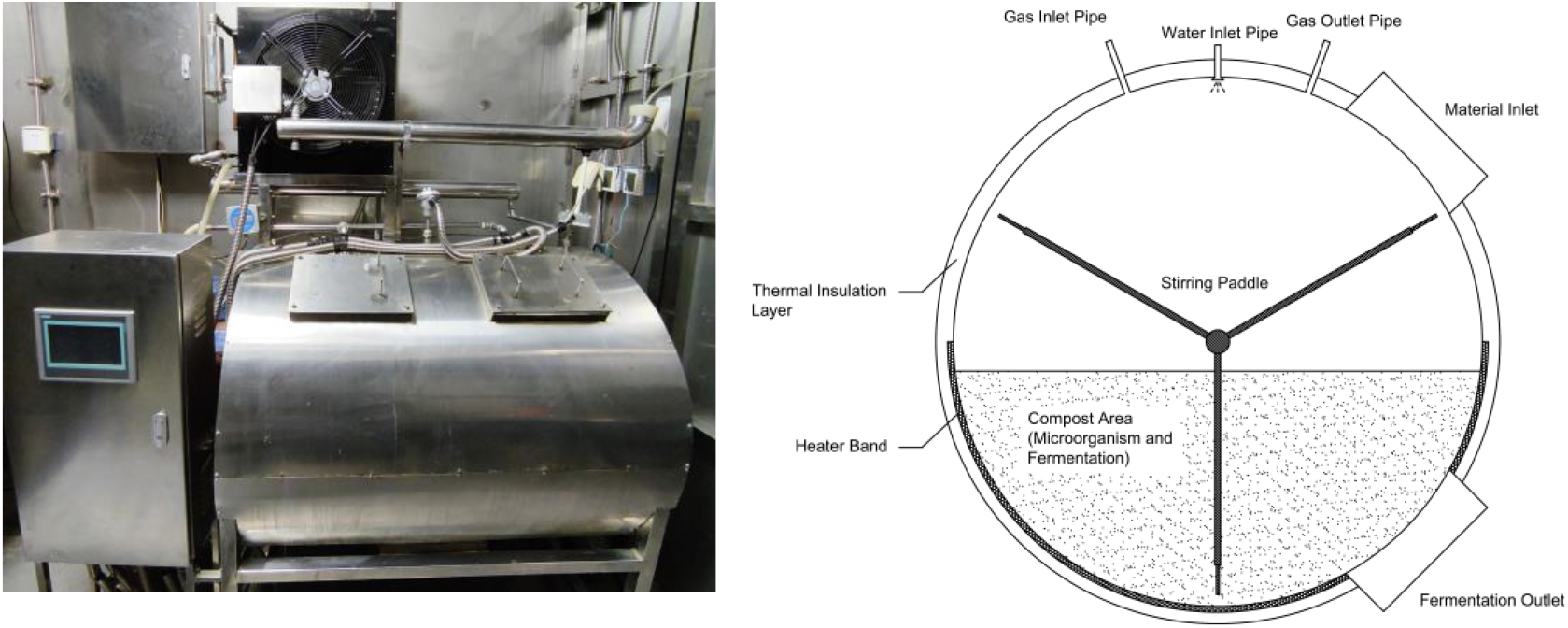
Prototype of SWTU (left) and internal structure diagram (right)

#### 2.3.1 Physical structure

The size of prototype of SWTU is 130×105×185cm. Its total volume is 460L with 150L maximum capacity of materials and maximum power consumption of 8500W.

The solid waste mainly includes straw and chaff of wheat *(Triticum aestivum* L.), frass of yellow mealworm (*Tenebrio molitor* L.) and feces of crew produced during the 370-day experiment in LP1. The raw materials of straw and chaff are crushed into fragments of 0.5–1 cm with a disintegrator (DF-20, Wenling LINDA Machinery Co. Ltd., China) before being put into the SWTU continuously. The microbial inoculants for solid waste fermentation are selected according to the previous experiments in our group (He et al., 2010).

#### 2.3.2 Operation technology

The number of microorganisms is counted by spread plate method.

The contents of acid-insoluble ash are determined according to Van Soest’s method (Strayer et al., 1997; Van Soest et al., 1991), using the crude fiber tester FIWE six raw fiber extractor (VelpScientifica, Italy).

The CO_2_ concentrations at the air inlet and exhaust are detected by online CO_2_ sensors (MH-Z14NDIR, scale: 0–10,000 ppm). The high percentage of O_2_ in the air and its small variation make it impossible for sensors to accurately measure the small change of O_2_ concentration. The O_2_ consumption needs to be replenished by interval ventilation during the operation of the system. The ventilation rate is calculated based on the percentage of air pump working time, and the duty cycle can be set according to the actual demand. Constant-flow air pump is applied for ventilation at a speed of 16.5L/min to keep the ventilation rate in the range of 5-20%. Exhaust gas is air-cooled for the recovery of water vapor in the exhaust gas, which will condense into the water storage tank for material humidification. The remaining will be sterilized by ultraviolet and adsorbed by activated carbon to kill the microorganisms escaping with the exhaust gas and adsorb harmful organic volatiles and odor gases to prevent pollution of the cabin atmosphere.

The temperature sensors are used to monitor the heating sheet temperature and material temperature. The appropriate range for fermentation, 40-60°C, is determined from our previous study(G. Liu et al., 2016), which regulates the rate of fermentation. To control the temperature, resistance wires are used for heating and glass wool is used for insulation.

The humidity sensor can monitor the moisture content of the material and the partial pressure of the vapor. When the system starts, the pump delivers water through the supply line, then the atomization nozzle atomizes and sprays it out to control the humidity in the range of 50-70%.

The material is completely mixed by motor-driven paddles intermittently once an hour.

### 2.4 Parameter identification and model validation

#### 2.4.1 Preprocessing of experimental data

After the 370-day experiment, vast amounts of experimental data on SWTU operation were collected for parameter identification and model validation. First of all, experimental data were preprocessed by the following steps:

1. Removal of outliers: Data falling outside the range of the sample mean ± 2.5 times standard deviation are considered outliers and abandoned.
2. Removal of high-frequency noise: In sensor-based sampling, the recorded signals are inevitably mixed with high-frequency noise, which should also be removed before parameter identification and validity check to ensure accuracy. In this study, a Butterworth low-pass filter is designed to remove high-frequency noise, in which the order of the filter is 4 and the normalized frequency is 0.65.
3. Removal of the linearized trend of data: By doing that, the analysis can be focused on the main nonlinear fluctuating trend of data, which allows the data to better reflect the dynamic characteristics of actual SWTU.
4. Interpolation to generate missing data: In the sampling process, some data are missing. Because of the short sampling interval, interpolations have to be used to predict the values of missing data, so that sufficient data are available for parameter identification and model validation. In this study, B-spline function interpolation is used to predict value at each time point with missing data.

After data preprocessing, the time-series experimental data were divided into two parts, half of them were applied for parameter identification and the other half for model validation.

#### 2.4.2 Methods for parameter identification and model validation

The simulation options have been properly set according to the complexity, accuracy, speed, and the computational expense of the digital simulation for parameter identification and model validation (Table 1).

**Table 1.**
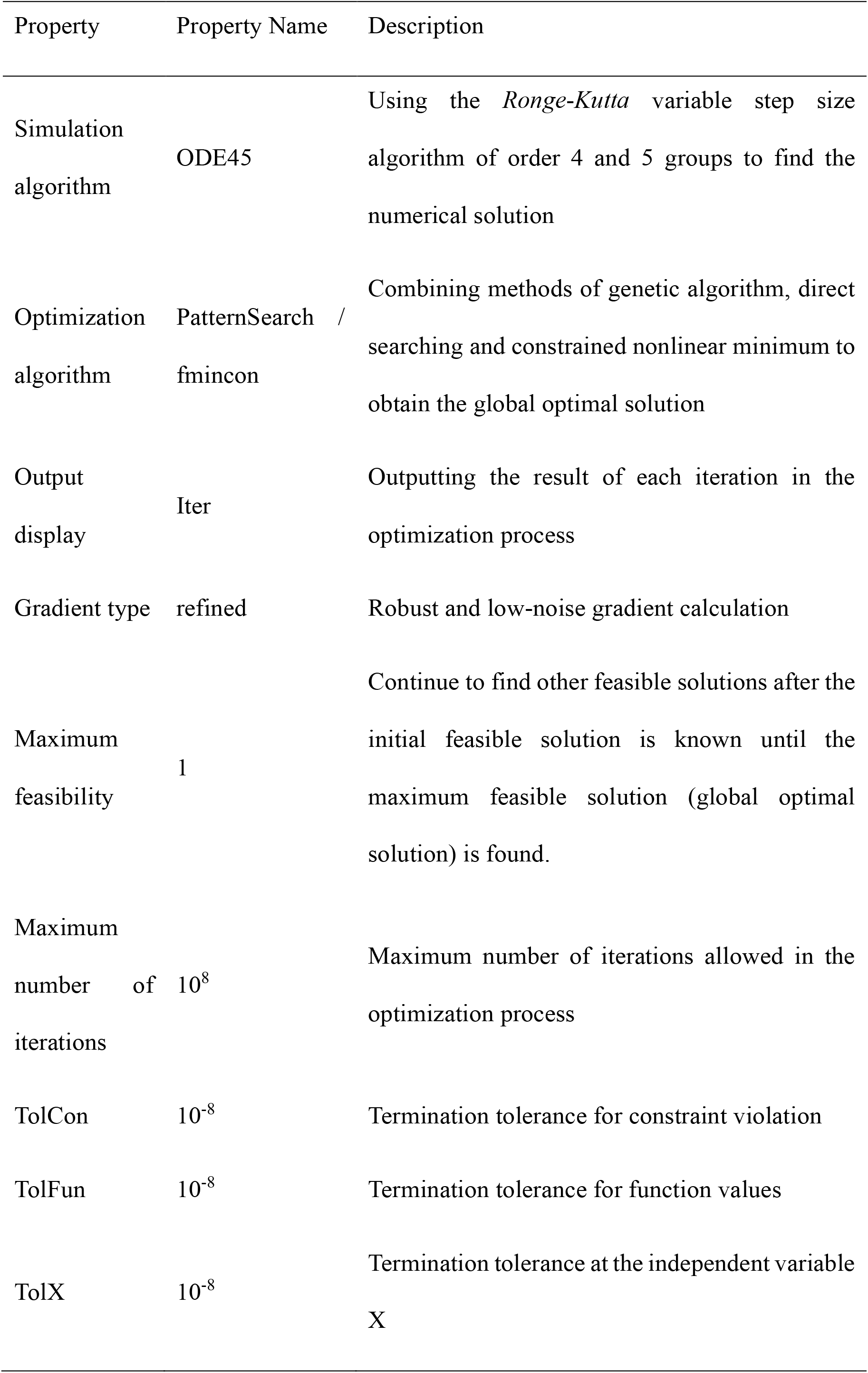
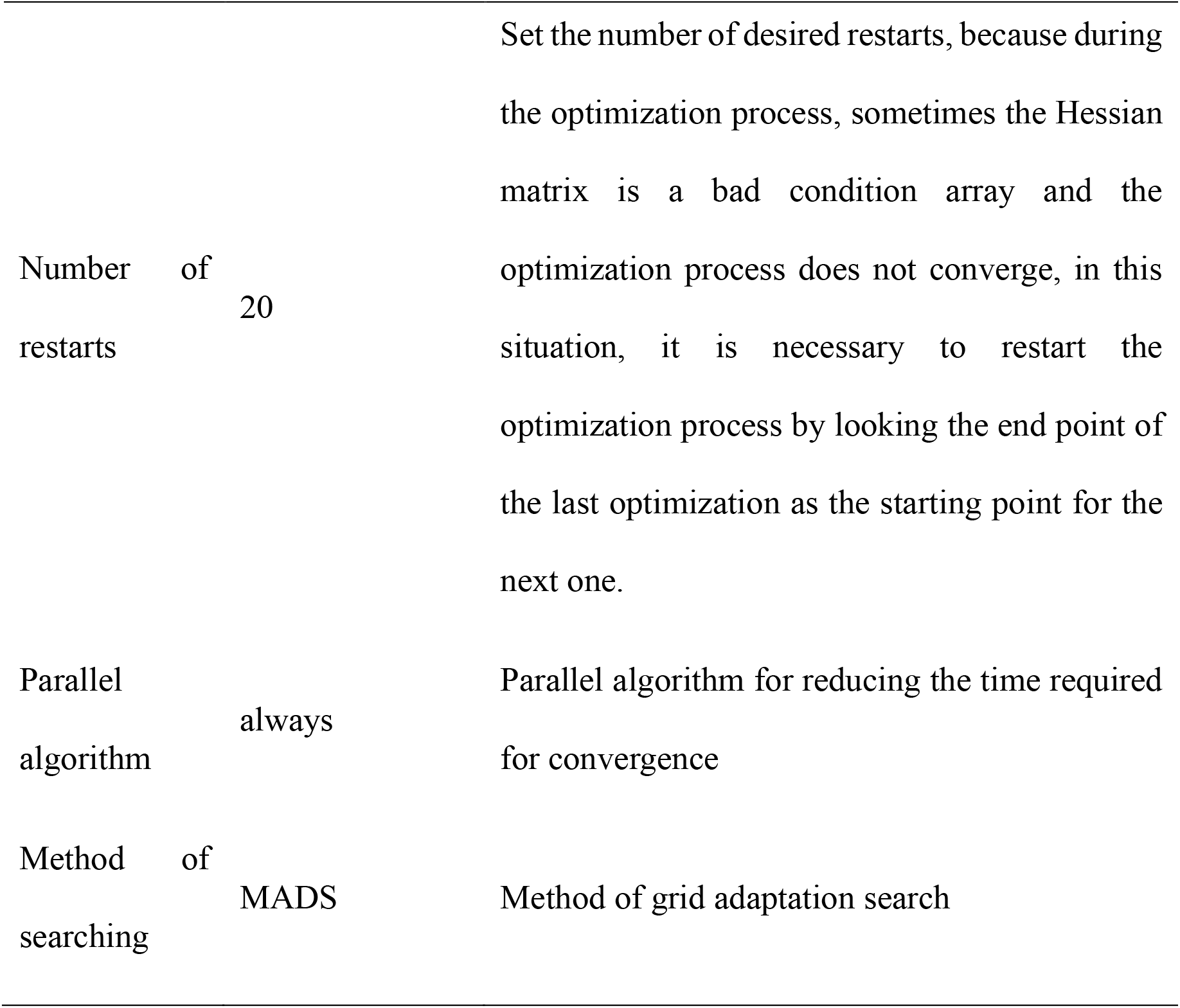
Options set for parameter identification and model validation.

| Property | Property Name | Description |
| --- | --- | --- |
| Simulation<br>algorithm | ODE45 | Using the *Ronge-Kutta* variable step size algorithm of order 4 and 5 groups to find the numerical solution |
| Optimization<br>algorithm | PatternSearch /<br>fmincon | Combining methods of genetic algorithm, direct searching and constrained nonlinear minimum to obtain the global optimal solution |
| Output<br>display | Iter | Outputting the result of each iteration in the optimization process |
| Gradient type | refined | Robust and low-noise gradient calculation |
| Maximum<br>feasibility | 1 | Continue to find other feasible solutions after the initial feasible solution is known until the maximum feasible solution (global optimal solution) is found. |
| Maximum<br>number of<br>iterations | $10^8$ | Maximum number of iterations allowed in the optimization process |
| TolCon | $10^{-8}$ | Termination tolerance for constraint violation |
| TolFun | $10^{-8}$ | Termination tolerance for function values |
| TolX | $10^{-8}$ | Termination tolerance at the independent variable X |
| Number of restarts | 20 | Set the number of desired restarts, because during the optimization process, sometimes the Hessian matrix is a bad condition array and the optimization process does not converge, in this situation, it is necessary to restart the optimization process by looking the end point of the last optimization as the starting point for the next one. |
| Parallel algorithm | always | Parallel algorithm for reducing the time required for convergence |
| Method of searching | MADS | Method of grid adaptation search |

## 3. Results and discussion

### 3.1 Parameter identification

The 370-day experiment in LP1 showed that SWTU prototype treated 160kg (dry weight) higher plants inedible biomass, 56kg (dry weight) feces, consumed 587L water and produced 68kg (dry weight) soil-like substrate used for higher plants cultivation, and thus satisfactorily fulfilled the task of solid waste treatment.

The parameter identification was conducted by experimental data and digital simulations, and the theoretical values of parameters were used for initial values. During the digital simulation, the parameter identification process went on smoothly with fast convergence (Table 2). Yielded parameters resulted that predicting values were very close to experimental data.

**Table 2.**
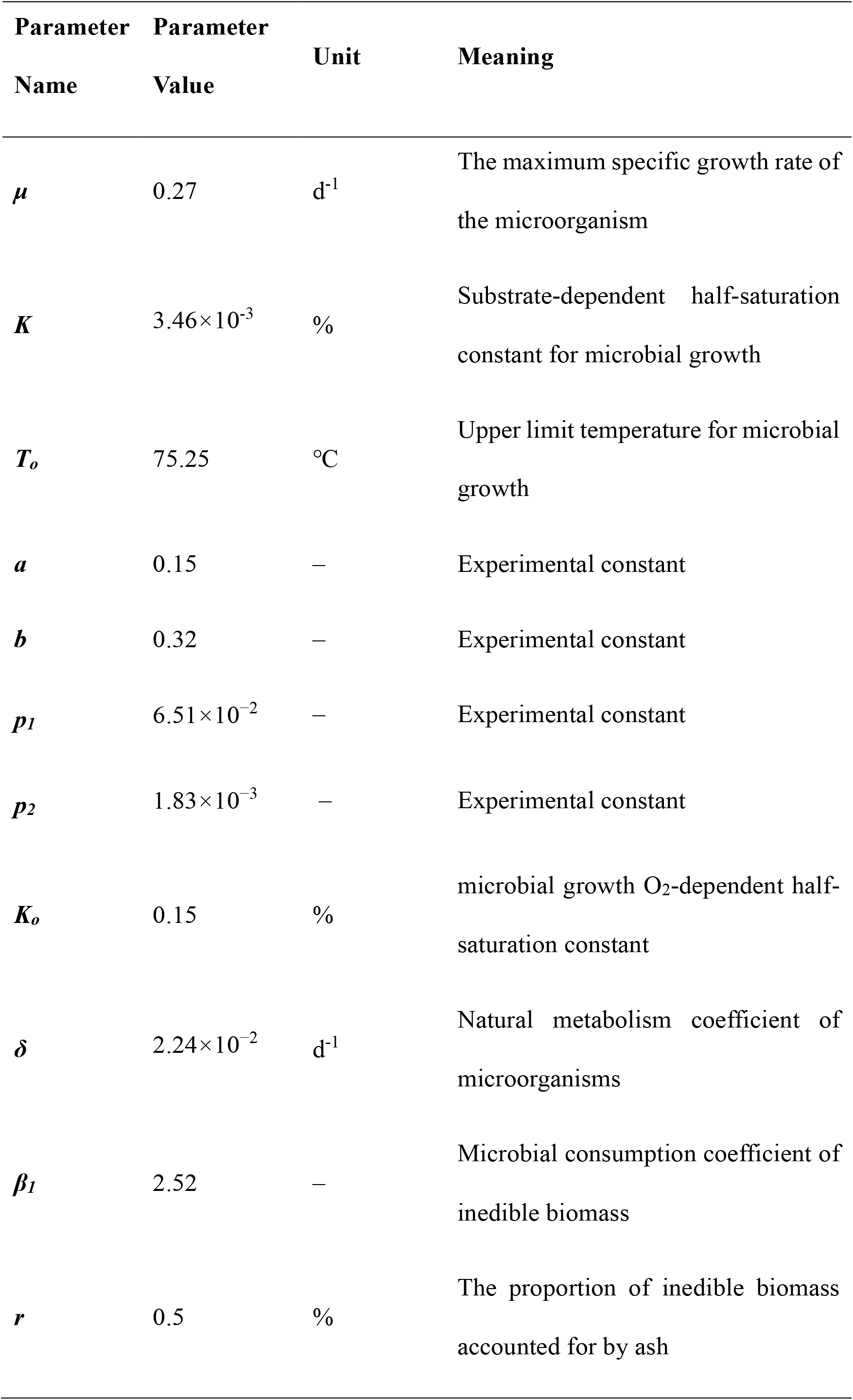
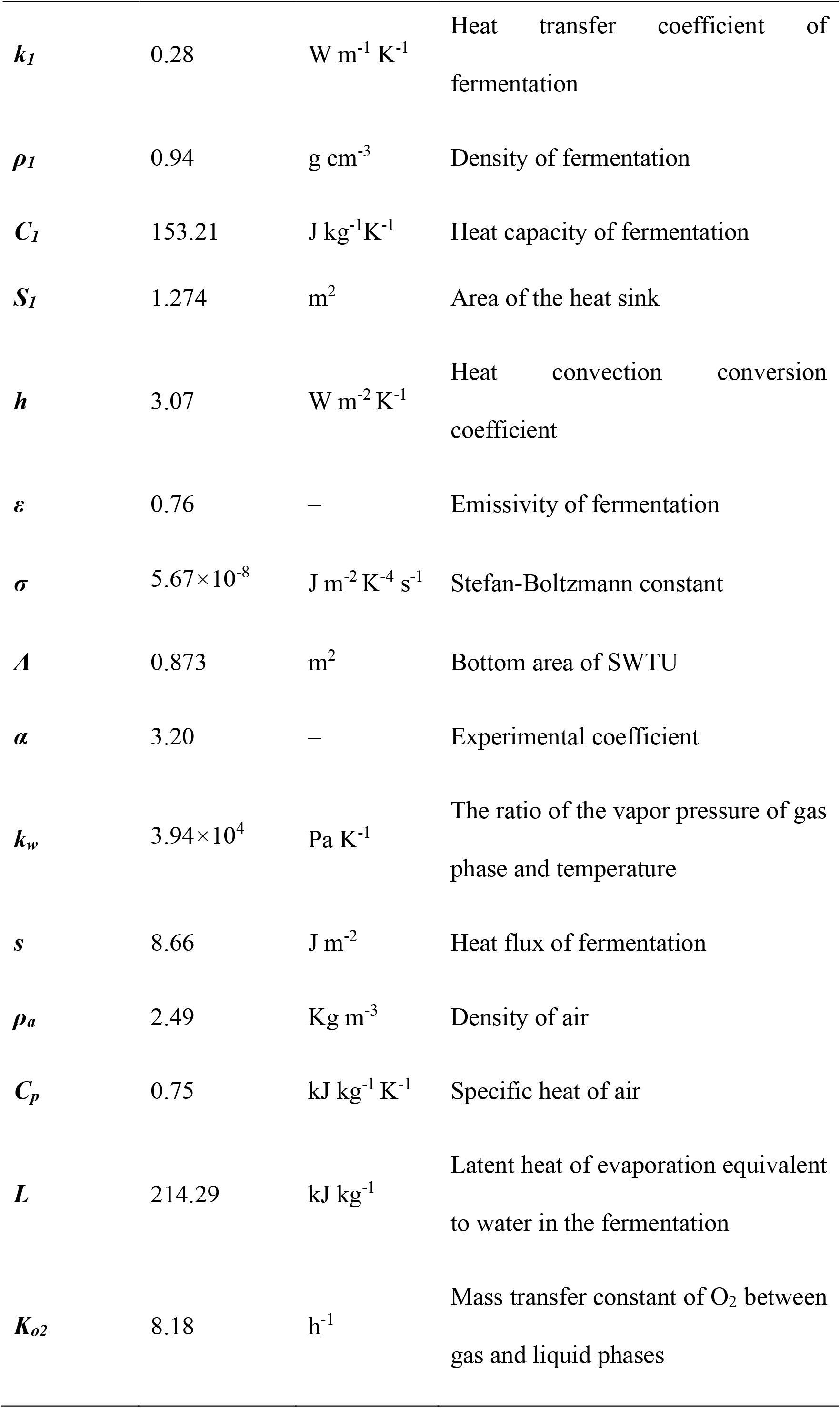
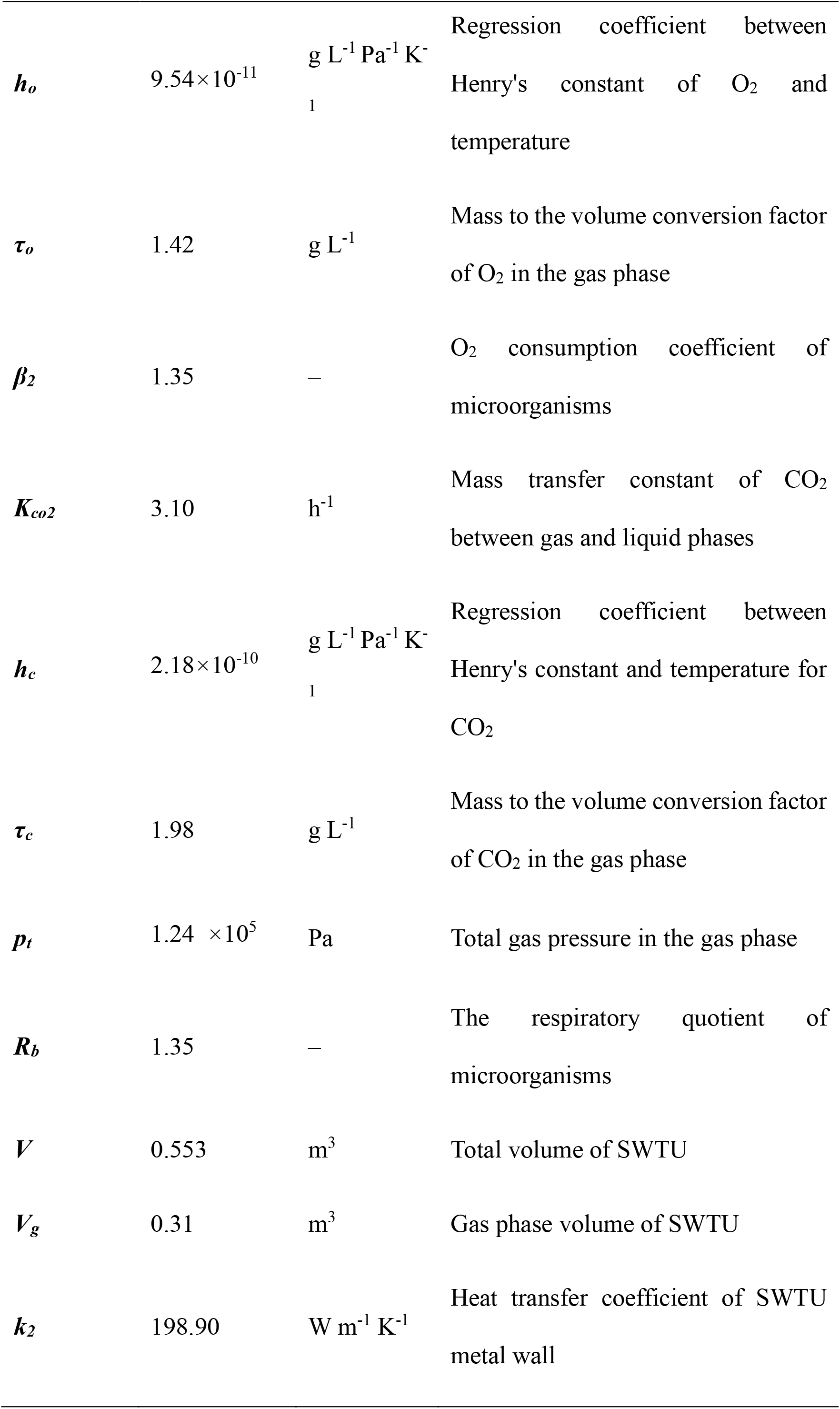
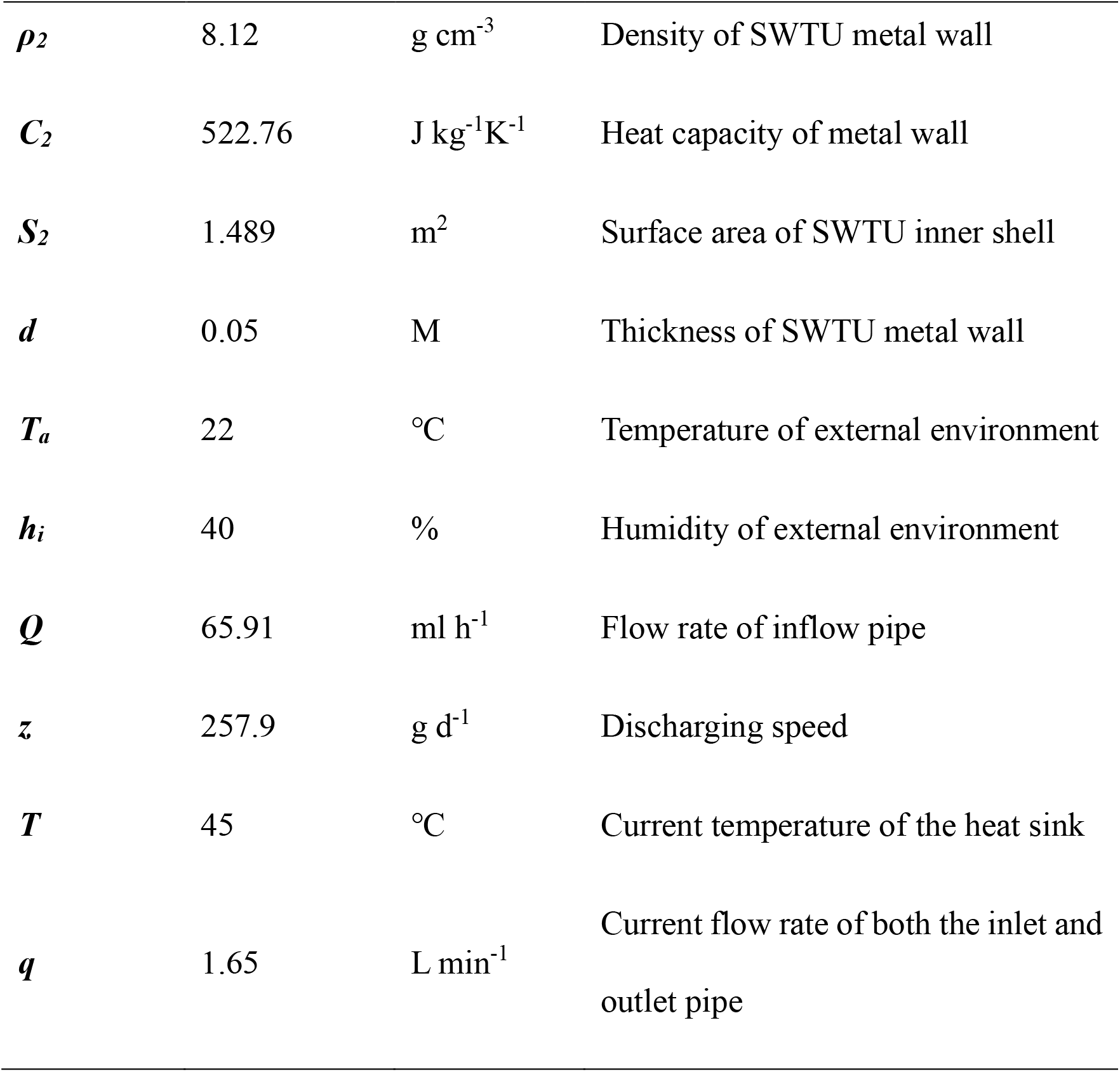
Result of parameter identification of SWTU kinetic model.

Some parameters have been specified, and need not to be identified by experiment, such as areas of the heat sink, bottom area of SWTU, total volume of SWTU, gas phase volume of SWTU, the surface area of SWTU inner shell, the thickness of SWTU metal wall, the temperature of the external environment, the humidity of the external environment.

### 3.2 Model Validation

Based on identified parameters and the left experimental data, the SWTU simulation is performed again, and then the predicted values are compared with the remaining half of the experimental data to test the behavioral similarity between the prototype and the model of SWTU. As illustrated in Figure 5, *r* denotes Spearman rank correlation coefficient, *NSE* is Nash-Sutcliffe efficiency standard deviation between simulating data (*ŷ*) and experimental data (y). The *NSE* is defined as follows:

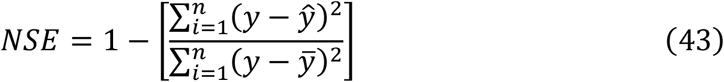

In terms of the model validation, the smaller the *NSE*, the higher the model validation. The *F*-value represents statistical regression effects of SWTU kinetic model. The significance probability p of Spearman rank correlation coefficient and model mechanism explanation effects < 0.01. Therefore, the conclusion could be drawn that the kinetic model of SWTU is exceedingly similar to its prototype in dynamic behaviors.

**Fig. 5:**
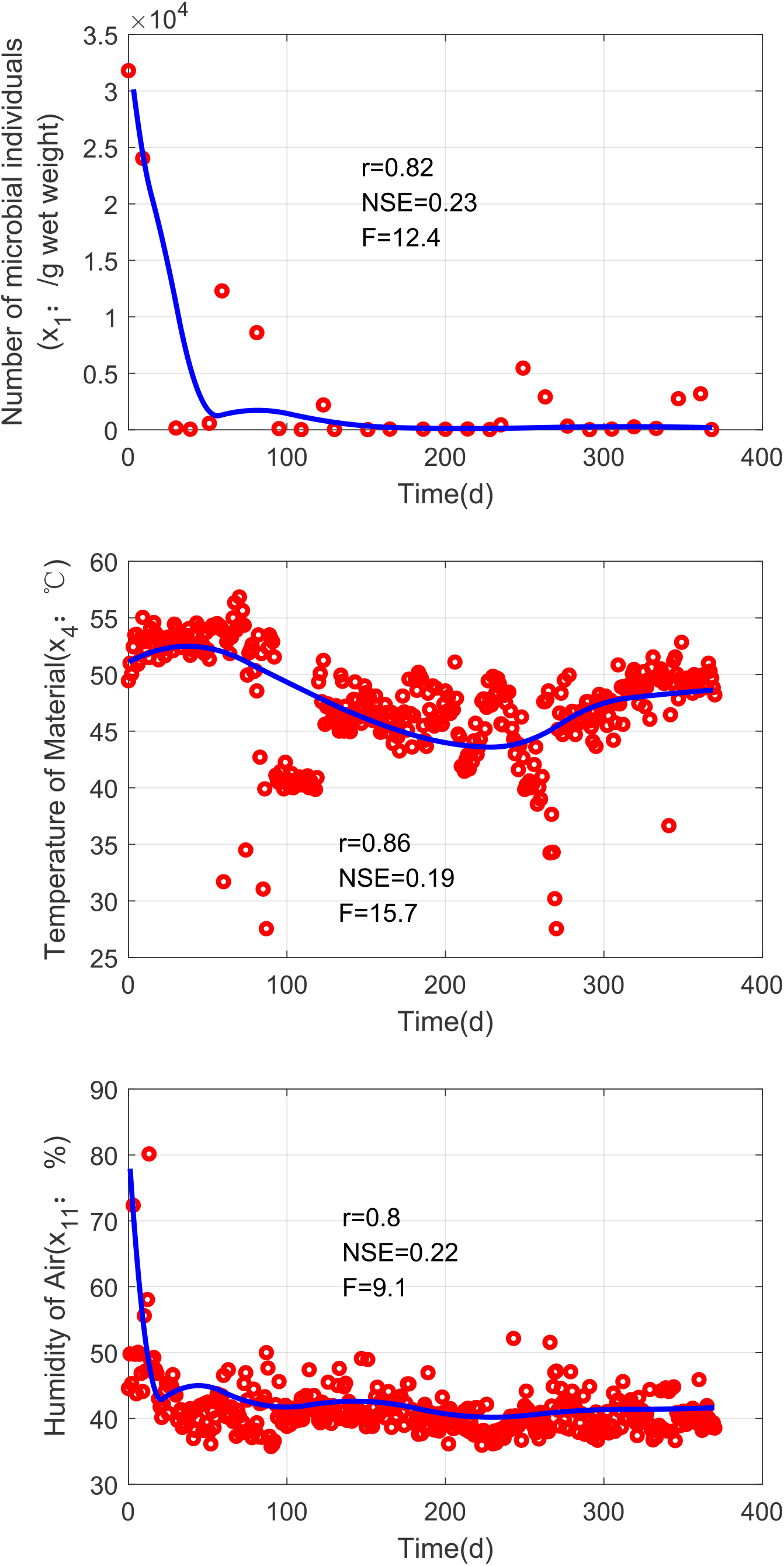
Validity test of SWTU model.

### 3.3 Discussion

#### 3.3.1 Structural similarity between model and prototype of SWTU

Since the behavior similarity between the model and prototype of SWTU has been validated by comparing simulated values with experimental data, the structural similarity could also be accredited as follows:

Firstly, the SWTU prototype is designed and built from its upfront kinetic model directly and has successfully maintained a long-term operation with robust stability and desired dynamic performance in LP1. At the same time, the experimental data can sufficiently give rise to the rapid convergence of the parameter identification process. Both signified that there is structural similarity between the kinetic model and SWTU prototype (Fig. 1). In a word, this kinetic model has accurately described the interactions and relationships between SWD processes and biotic/abiotic environmental factors, which are effective dynamic mechanisms driving the operation of SWTU.

Secondly, although the existing models focused on SWD processes affected by environmental factors(Ahn et al., 2008; Białobrzewski et al., 2015a; Rafiee et al., 2017; Solemauri et al., 2007; Vasiliadou et al., 2015a), the kinetic model in this study differs in the form, in order to effectively guide and serve the engineering application. For example, the temperature is a key factor to influence the SWD processes because the growth and metabolites of microorganisms are exquisitely sensitive to temperature change. The temperature influence functions are often pretty complicated to elaborate upon the detailed mechanisms(Białobrzewski et al., 2015b; Rosso et a l., 1995; Vasiliadou et al., 2015b). However, digital simulations of these kinetic models must take a long time, which is disadvantageous to engineering application. The temperature function proposed in (Eq.(3)) is also a bell-like function, which could greatly simplify the formula and calculation without diminishing the validity. Simultaneously, the other rate equations are also substantially simplified and tightly coupled together. More significantly, the microbial growth rate playing a fundamental role need to be modelled very precisely, while the other rates are specified on basis of it through coefficients calibration. This necessary simplification can enhance computational efficiency of digital simulation considerably to better serve model-based prototype design, building operation, real-time control and optimization of SWTU. As is known, the mathematical models of living systems are empirical or semi-empirical, there are no absolutely constitutive relationships like Newton’s law or Maxwell equations in physics to quantitively express the dynamic characteristics of living systems. Nevertheless, it does not affect the utility of models to understand natural principles and put them into engineering application. Although this kinetic model is not the ultimate version to quantitatively described the SWTU in contrast to other kinetic models, a great number of digital simulations did not give any self-contradictory results. Therefore, it is sufficiently self-consistent, and could be applied for mechanism explanation and engineering application.

#### 3.3.2 Further application in ecological engineering

Despite the kinetic model in this study is currently just the blueprint for design, building and operation of SWTU prototype, it could also be applied for further analysis, synthesis, control, and optimization of SWTU. For example, appropriate tunable parameters (such as temperature, humidity, discharging speed, flow rate of the inlet/outlet pipe) and proper feedback signals (such as microbial biomass, CO_2_concentration, O_2_ concentration) could be selected for closed-loop control of operation. Based on dissipative structure theory, a desired feedback control law for highly-nonlinear SWTU would be obtained from a Lyapunov function with specific ecological and engineering significance (Unpublished Work). According to cybernetics, the closed-loop SWTU control system can effectively resist different internal variations and external disturbances to keep robust operation with satisfying dynamic performance.

Furthermore, this kinetic model can also be extended to comprehend the dynamic mechanisms of physiochemical and ecological processes occurring in terrestrial (Jafarpour & Khatami, 2021) and riverine ecosystems (Koley, 2021) via appropriate model calibrations. Based on calibrated models and tunable parameters, relevant policies of environmental management and technologies of ecological engineering could be developed to promote sustainable development of the regional economy, society and environment.

## 4. Conclusion

In this study, a kinetic model of SWTU was upfront developed based on microbial ecology, system dynamics, cybernetics and digital simulation. The values of state variables sufficiently represented the components and functions of SWTU, and the rate equations accurately described the relationships and interactions between SWD processes and biotic/abiotic factors. Then a specific SWTU prototype was designed and built from this model and implemented during a 370-day experiment in LP1, the result of parameter identification and model validation through experimental data sufficiently demonstrated that the kinetic model of SWTU was greatly valid due to high structural and behavioral similarity with its prototype. Based on the kinetic model, further researches on closed-loop control and process optimization could be carried out for the construction of an advanced SWTU prototype. The kinetic model and the applied method of model-based prototype design, building and optimization can be used for other purposes, such as environmental management, ecological restoration, sustainable development.

## Abbreviation

SWD: solid waste decomposition
SWTU: solid waste treatment unit
LP1: Lunar Palace 1
BLSS: Bioregenerative Life Support System

